## Supplementary file for "Structure of the human systemic RNAi defective transmembrane protein 1 (hSIDT1) reveals the conformational flexibility of its lipid binding domain"

### Supplementary figures

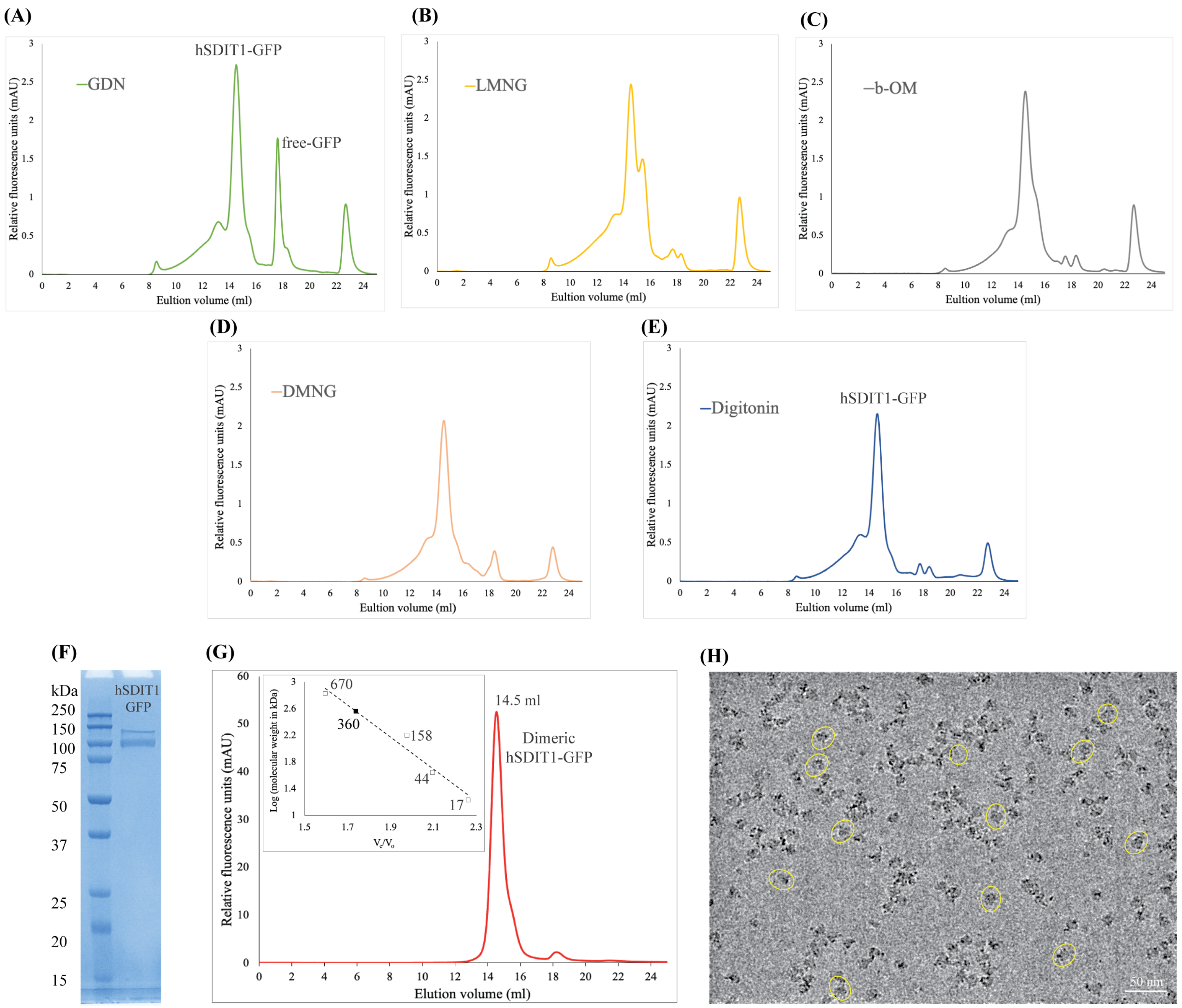


**Figure S1: hSIDT1 purification and cryo-EM sample preparation**

**(A-E)** Comparison of the relative solubility and homogeneity in 1% of GDN, LMNG, β-OM, DMNG, and digitonin, respectively. All the detergents were prepared in 25 mM Tris-HCl, pH 7.5, 200 mM NaCl, 1 mM PMSF, 0.8 µM aprotinin, 2 µg/mL leupeptin, and 2 µM pepstatin A. **(F)** SDS-PAGE (12%) showing the purity of hSIDT1-

GFP. Although the estimated monomeric molecular weight of the fusion protein is ~ 124 kDa, like other eukaryotic membrane proteins, hSIDT1 displays anomalous electrophoretic mobility. **(G)** Intrinsic tryptophan fluorescence of purified hSIDT1-GFP was monitored in a size-exclusion chromatography experiment on Superose 6 Increase 10/300 GL column at 0.5 ml/min flowrate. Black square on the standard plot (log of protein molecular weight (kDa) vs. ratio of the elution volume to the void volume (Ve/Vo) indicates that recombinant hSIDT1-GFP fusion, which elutes at 14.5 ml corresponding to a dimer molecular weight of ~360 kDa (hSIDT1-GFP dimer + digitonin micelle). **(H)** Representative micrograph showing vitrified hSIDT1-GFP (yellow circles) at 1 mg/ml on UltrAuFoil holey-gold 300 mesh 1.2/1.3 µm grids, imaged using Titan Krios G4i transmission electron microscope. The protein distribution on the grids, SDS-PAGE, and size-exclusion chromatogram indicate a monodisperse sample preparation. Scale bar is 50 nm.

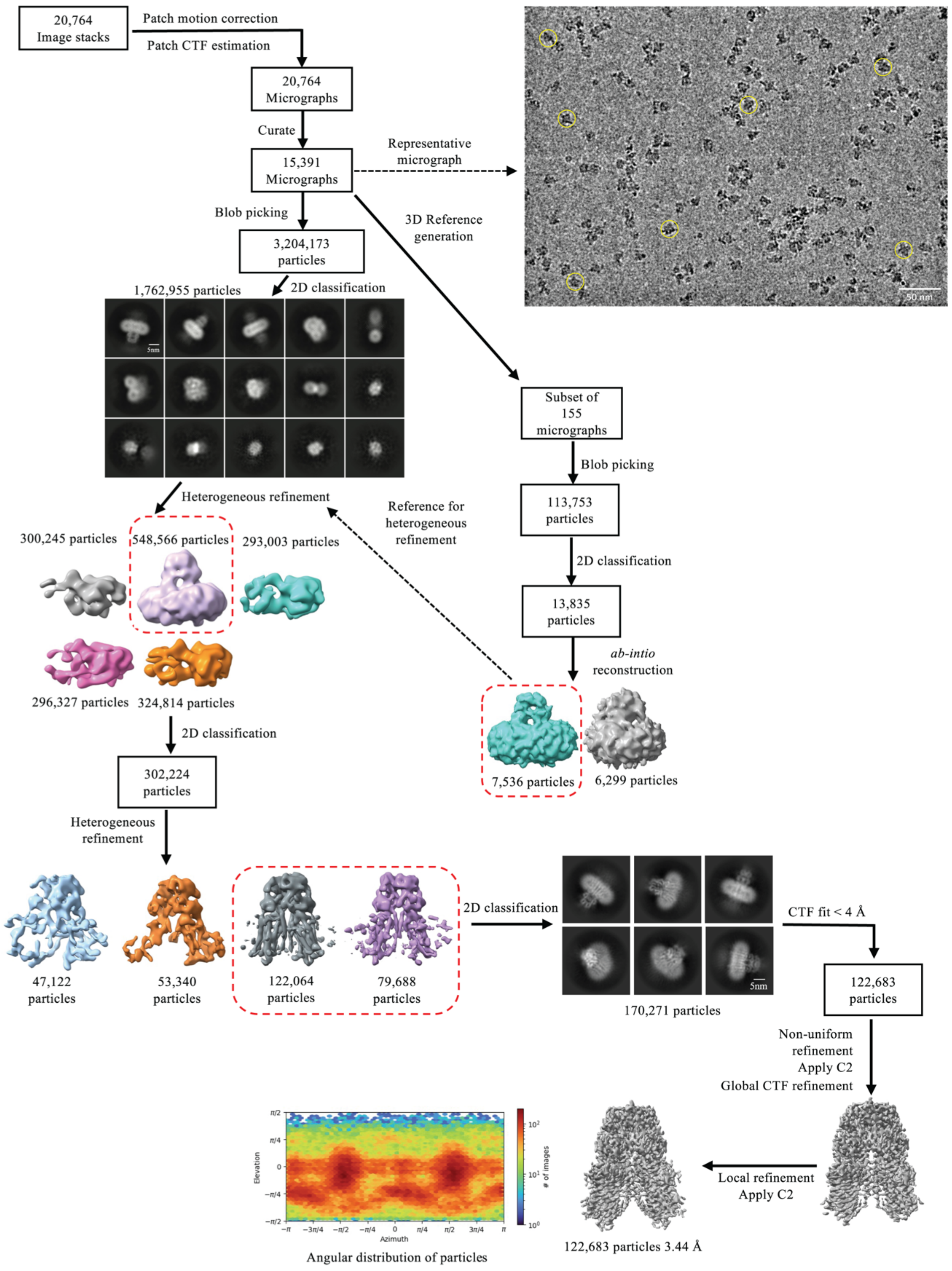


**Figure S2: Data processing workflow of hSIDT1-GFP cryo-EM data**

The data was processed using CryoSPARC (v 4.2.1) [52]. On the top right, a representative motion-corrected micrograph with hSIDT1-GFP particles (yellow circles) is highlighted. A subset of 155 motion-corrected micrographs was selected, and particles were picked by blob picker and cleaned by iterative 2D classification to generate a low-resolution *ab initio* reconstruction to be used as a reference input for heterogeneous refinement of the entire dataset (dashed box and arrow). Briefly, ~3.2 million particles were picked by blob picker, cleaned by 2D classification, and sorted further by heterogeneous refinement. ~548,000 particles from the best class (dashed box) in this heterogeneous refinement were further cleaned up by iterative 2D classification. ~302,000 particles were re-extracted unbinned in a 360 px box and sorted further by iterative heterogeneous refinement and 2D classification jobs. The resultant set of 170,000 clean particles was re-extracted from micrographs with a CTF fit <4 Å to yield a final stack of about 122,000 particles. C2 symmetry was applied at this stage, and non-uniform and CTF refinements were performed to obtain a final map at 3.44 Å [53]. The sharpened and unsharpened maps from this job were used for model building and analyzing the structure of hSIDT1-GFP. The final 3D model was validated using the sharpened map for deposition.

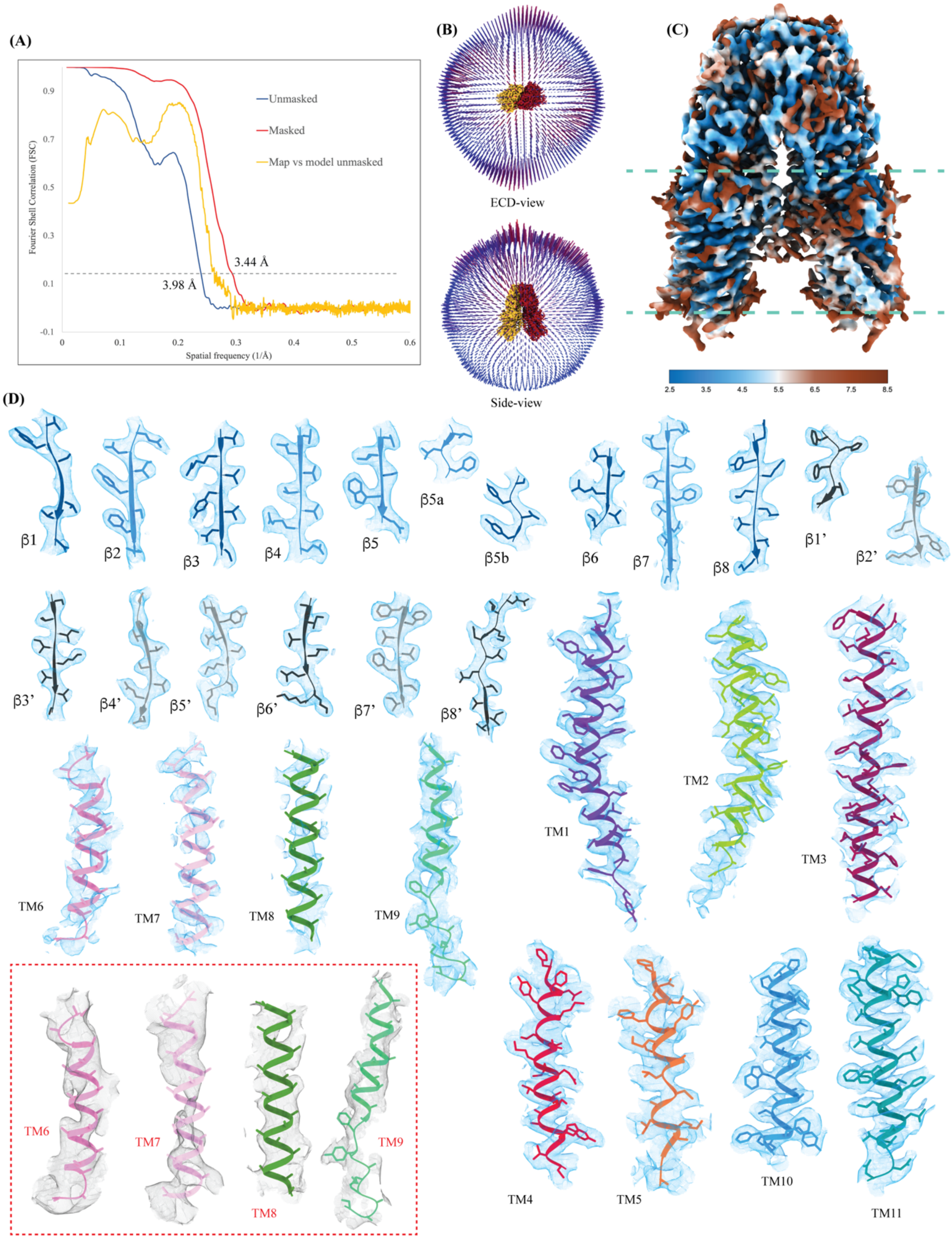


**Figure S3: Cryo-EM map quality and model building**

**(A)** The cross-validation Fourier shell correlation (FSC) curves for the final masked hSIDT1-GFP masked (red) and unmasked (blue) refinement maps and the model vs. final unmasked map (yellow). The gray dashed line indicates the $FSC_{0.143}$ threshold. FSC curves were calculated using Phenix's Mtriage [59]. **(B)** The angular distribution of the particles used in the final reconstruction in ECD- and side-view. **(C)** Estimated local resolution of the final C2 refined map (isosurface threshold level of 0.14 in ChimeraX). **(D)** The cryo-EM density of all the secondary structure elements is shown in light blue at a isosurface threshold level of 0.25 for the ECD region and 0.1 for the TMD region in ChimeraX [60]. Side chains for almost all the elements could be modeled unambiguously into the density except for TMs 6-8 and the extra-cytosolic half of TM9. In these regions, the C$\alpha$ traces were placed using an unsharpened map (red dashed box), and the side chains were trimmed to C$\beta$. ECD was modeled into the density using ModelAngelo [55]. The AlphaFold model of hSIDT1 (UniProt ID: Q9NXL6) was used to build the TMD core, and TMs 5-9 were manually built using Coot [57, 58].

**(A)**

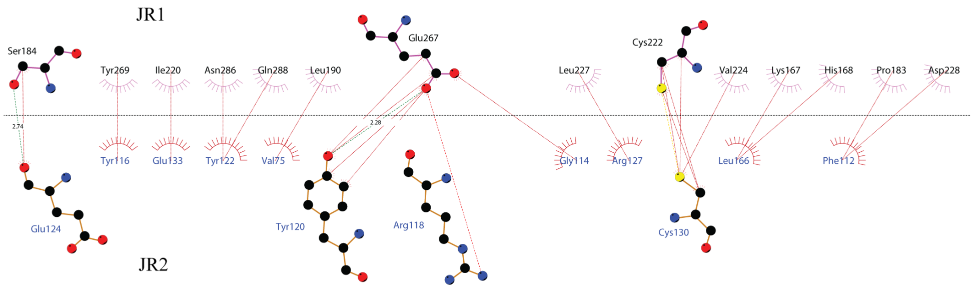


JR1-JR2 interface

**(B)**

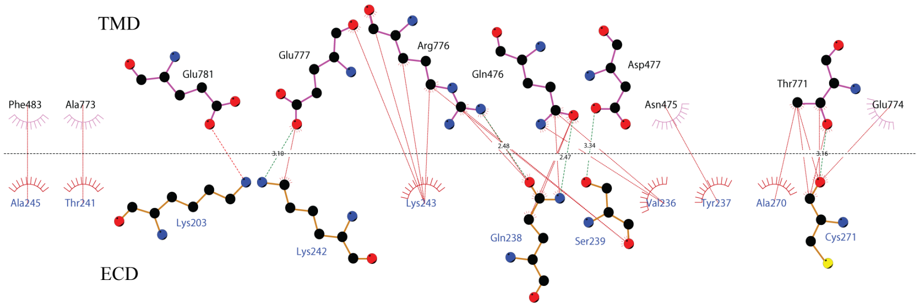


ECD-TMD interface

**(C)**

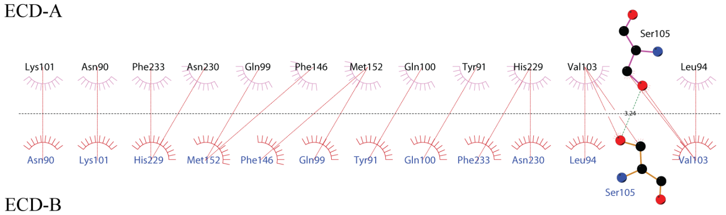


ECD-A-ECD-B interactions at the dimer interface (sub-interface 1 & 2)

**(D)**

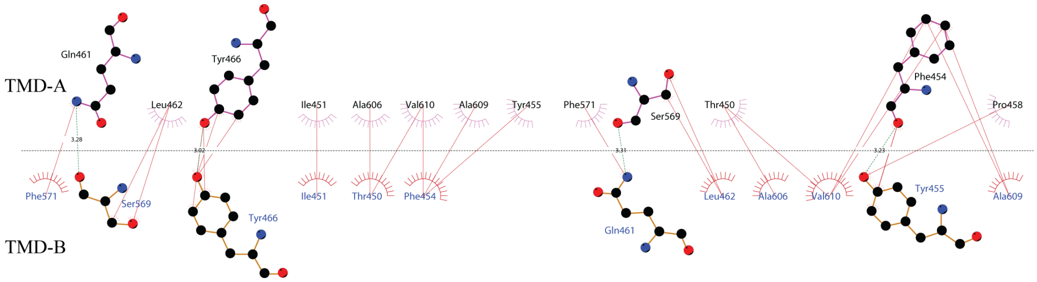


TMD-A-TMD-B interactions at the dimer interface (sub-interface 3)

**Figure S4: 2D plots of the critical inter-chain and intra-chain interactions**

LigPlot+ was used to generate the 2D depictions of the interactions (<4.5Å) at the **(A)** JR1-JR2 interface, **(B)** ECD-TMD interface, and **(C)** and **(D)** dimer interface [63]**.** Hydrogen bonds are depicted by dashed lines, critical residues are shown as ball & stick models, and amino acids involved in the non-bonded interactions are indicated as eyelashes. The sulfurs of disulfides are highlighted in yellow.

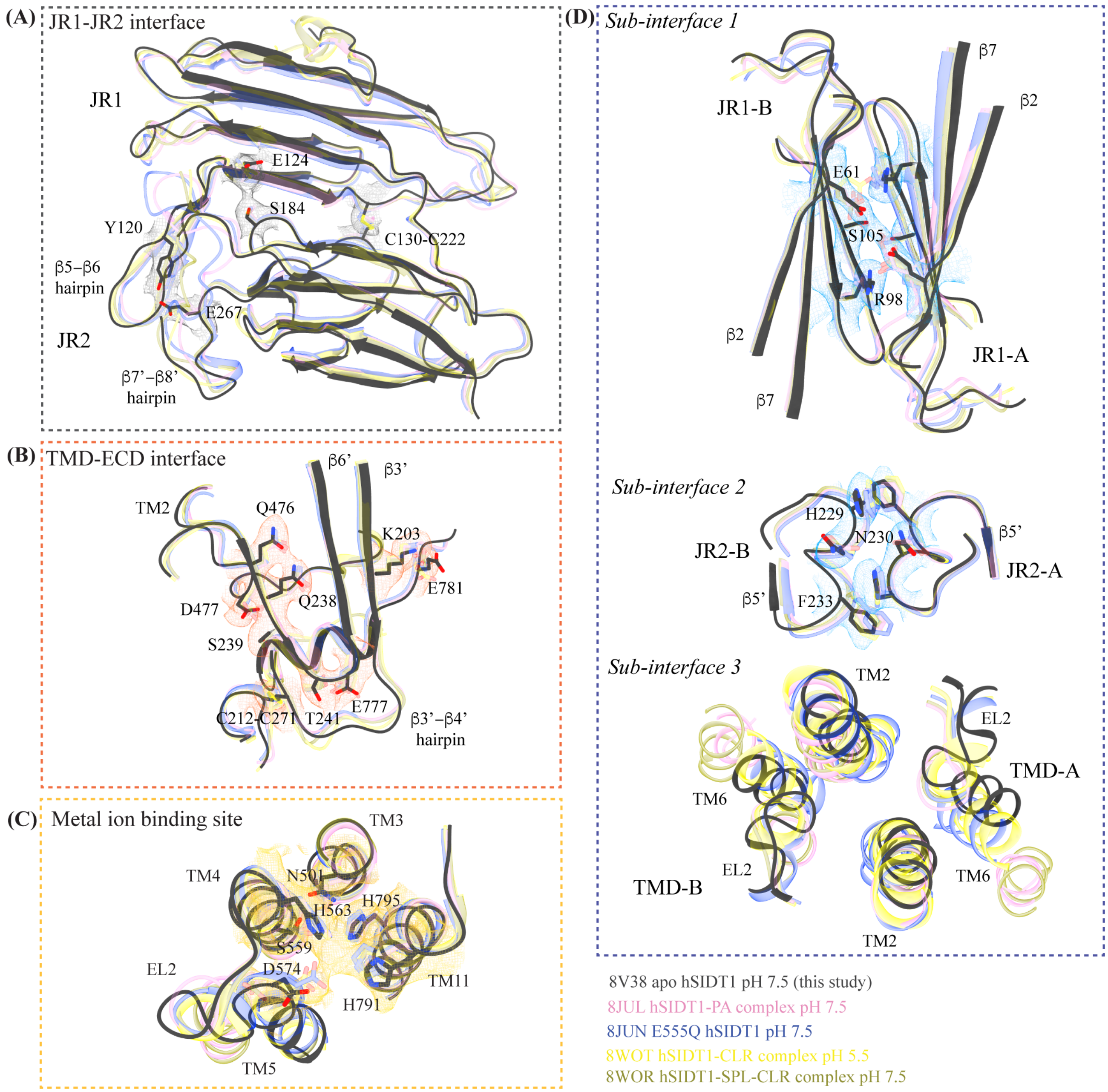


**Figure S5: Comparison of inter-chain and intra-chain interactions of different hSIDT1 structures**

Here we compared **(A)** JR1-JR2 interface (gray box), **(B)** TMD-ECD interface (orange box), **(C)** metal ion binding site (yellow box), and **(D)** dimer interface (blue box) of hSIDT1 structure obtained in different conditions

- 8V38 (apo hSIDT1 at pH 7.5; gray), 8JUL (hSIDT1-PA complex at pH 7.5; pink), 8JUN (E555Q-hSIDT1 at pH 7.5; blue), 8WOT (hSIDT1-CLR at pH 5.5; yellow), 8WOR (hSIDT1-SPL-CLR at pH 7.5; green). The amino acids involved in hydrogen bonding and ring stacking interactions at the interfaces are highlighted. The densities for these amino acids in 8V38 have been displayed in colors that match their insets (isosurface threshold level of 0.25 for the ECD region and 0.1 for the TMD region in ChimeraX). We notice that the differences in the elements that form these interfaces lie primarily within TMD at sub-interface 3.

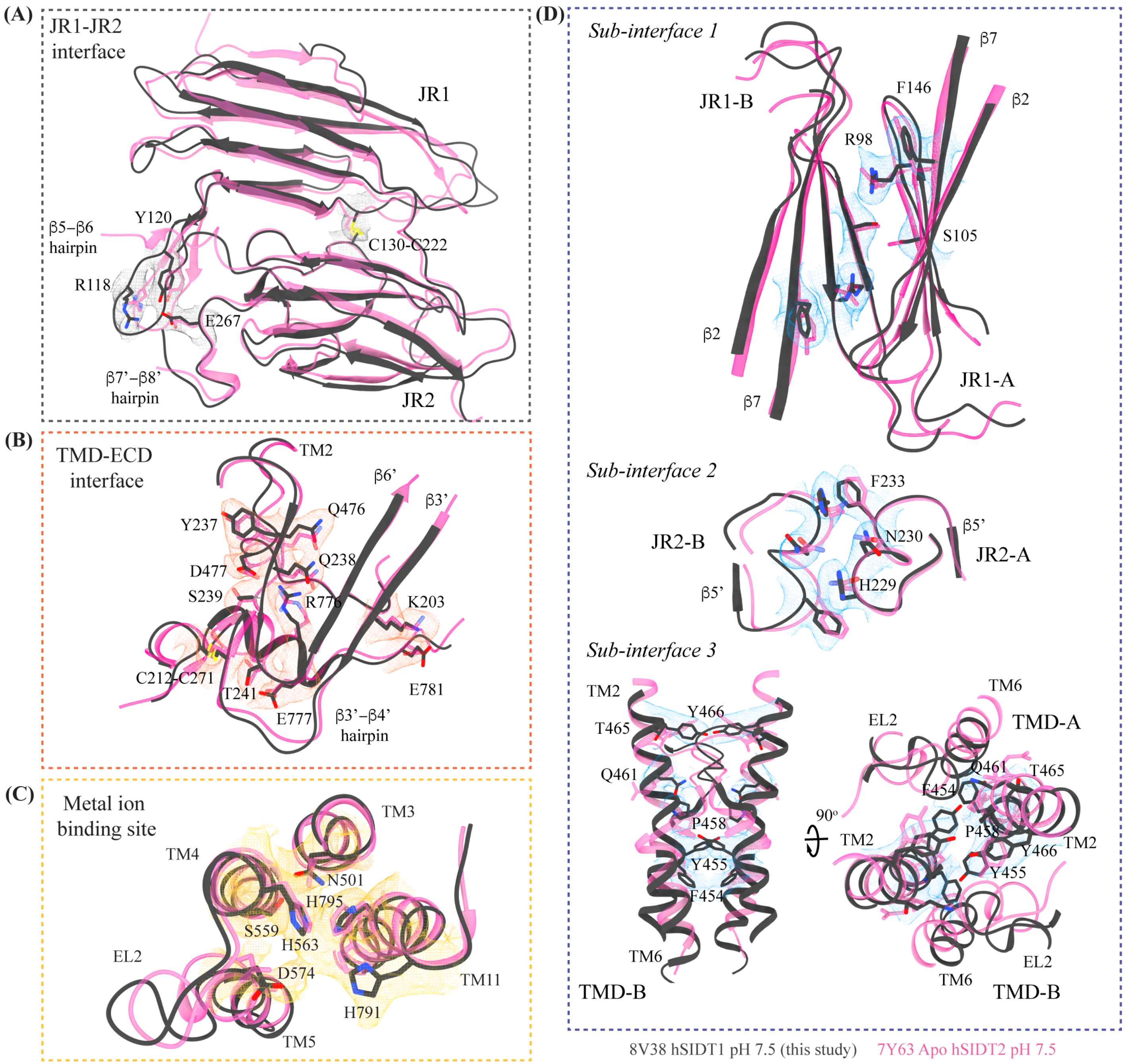


**Figure S6: Comparison of inter-chain and intra-chain interactions of hSIDT1 and hSIDT2**

Here we compared **(A)** JR1-JR2 interface (gray box), **(B)** TMD-ECD interface (orange box), **(C)** metal ion binding site (yellow box), and **(D)** dimer interface (blue box) of hSIDT1 (8V38: gray) and hSIDT2 (7Y63; dark pink). The densities for 8V38 at these interfaces have been displayed in colors that match their insets (isosurface threshold level of 0.25 for the ECD region and 0.1 for the TMD region in ChimeraX). We notice that the residues involved in hydrogen bonding and ring stacking interactions at these interfaces are conserved between hSIDT1 and hSIDT2, and their positions change only in the TMD at sub-interface 3.

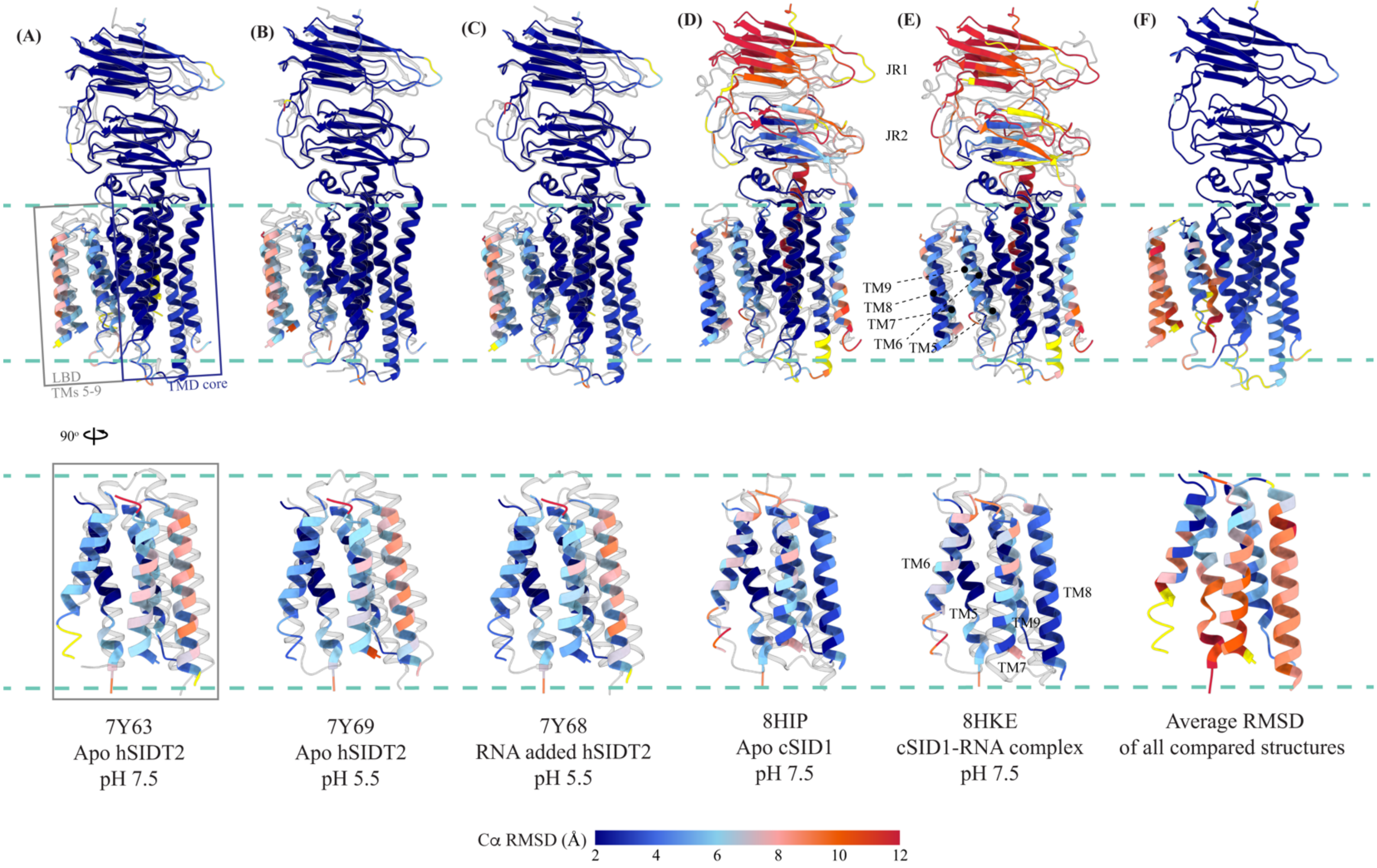


**Figure S7: Comparison of intra-chain dynamics of hSIDT1, hSIDT2, and cSID1 cryo-EM structures**

The panels **A-E** show chain A of apo hSIDT1 (8V38) colored by Cα RMSD (blue-to-red) obtained upon alignment with chain A of hSIDT2 or cSID1 structures. Yellow regions represent regions missing from the alignment. Bottom half of each panel shows only the LBD region. The structures compared with hSIDT1 in each panel are displayed in gray and the respective PDB IDs are indicated below each panel. Panel **F** represents chain A of 8V38 colored by average Cα RMSD obtained upon all-vs-all alignment with chain A of all the hSIDT1, hSIDT2 and cSID1 structures compared in this manuscript. Superpositions were performed using sequence based pairwise alignment in ChimeraX.

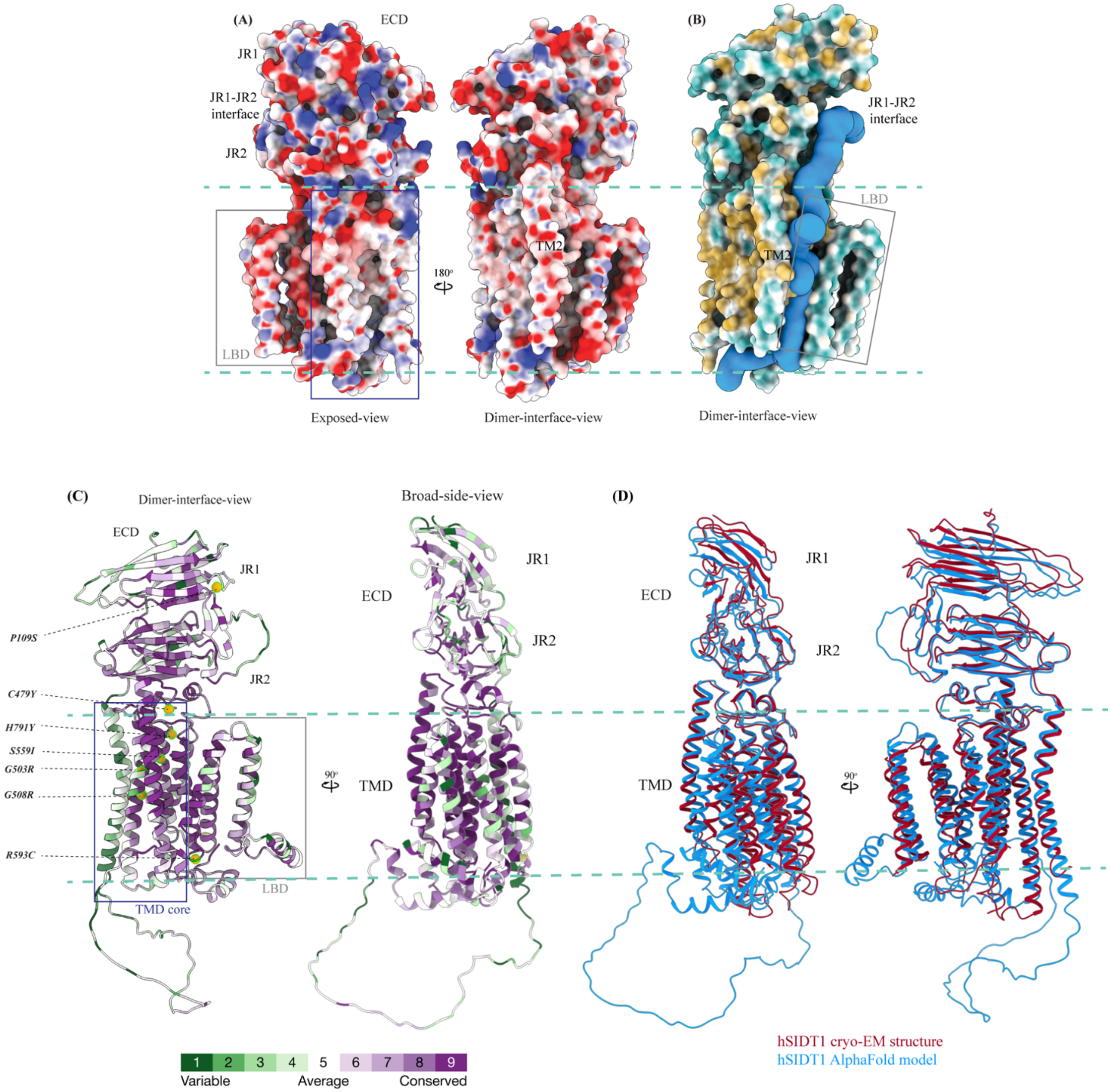


**Figure S8: Electrostatic potential, hydrophobicity, and sequence conservation of hSIDT1**

**(A)** Electrostatic potential and surface charge representation of hSIDT1 protomer. The display is contoured on the potential from -10 kT (red) to +10 kT (blue). Exposed surface of ECD, especially the JR1-JR2 interface shows a high density of basic residues conducive to RNA binding. The dimer interface of the TMD is lined with uncharged polar residues. **(B)** Surface representation of hSIDT1 protomer colored based on hydrophobic (orange) and hydrophilic (cyan) amino acids. In blue is the channel representing a potential path of RNA transport, which was predicted by MOLEonline server using a prove radius of 20 Å [61]. **(C)** hSIDT1 AlphaFold model (Q9NXL6) colored according to evolutionary sequence conservation scores generated by ConSurf webserver using homologs identified by PSI-BLAST and aligned by Clustal [64]. Equivalent mutations that attenuate systemic RNAi mediated by SID1 in *C. elegans* are highlighted as orange spheres. All of these mutations occur in the conserved regions of hSIDT1 and lie either in the path of the channel predicted by MOLEonline server or in the vicinity of the metal ion coordination site. **(D)** Superposition of hSIDT1 cryo-EM structure and AlphaFold model showing that the predicted differences lie primarily in JR1, and LBD regions of the proteins as observed in the comparisons using experimentally determined structures of hSIDT1, hSIDT2, and *C. elegans* SID1.

### Supplementary tables

### Table S1: Comparison of the sample preparation conditions employed by various studies

| | **Qian et al.,** [33] | **Zheng et al.,** [36] | **Hirano et al.,** [35] | **Sun et al.,** [34] | **Liu et al., [37]** | **This study** |
|---|---|---|---|---|---|---|
| | | **Expression** | | | | |
| **Cell type** | HEK293F | Sf9 | Expi293F | HEK293F | HEK293F | HEK293sGnTI- |
| **Expression construct** | Full-length hSIDT2 | ΔCL1-hSIDT1 and ΔCL1-hSIDT2 | Full-length hSIDT1 | Full-length hSIDT1 | Full-length hSIDT1 | Full-length hSIDT1-GFP |
| **Gene transfer** | Transient transfection | Baculoviral transduction | Baculoviral transduction | Transient transfection | Baculoviral transduction | Baculoviral transduction |
| **Cell culture time/temp** | 48 h/37 ºC | 60 h/27 ºC | 96 h/30ºC | 12 h/37 ºC<br>48 h/30 ºC | 12 h/37 ºC<br>36 h/30 ºC | 8 h/37 ºC<br>36 h/32 ºC |
| **Culture volume** | 12 L | 6 L | - | 1L | - | 1 L |
| | | **Purification** | | | | |
| **Cell lysis buffer** | 25 mM HEPES, pH 7.4, 150 mM NaCl, 1.95 μg/ml aprotinin, 1.5 μg/ml pepstatin, and 3 μg/ml leupeptin | 50 mM HEPES, pH 7.5, 300 mM NaCl, 1.04 mM AEBSF, 0.8 μM Aprotinin, 50 μM Bestatin, 15 μM E-64, 20 μM Leupeptin, 15 μM Pepstatin A, and 1 mM PMSF | 20 mM Tris-HCl, pH 7.0, 150 mM NaCl, and 1X protease inhibitor cocktail | 40 mM Tris-HCl, pH 7.5, 150 mM NaCl, 20% glycerol (v/v), and 1X protease inhibitor cocktail | 25 mM HEPES, pH 7.5, 250 mM NaCl, 5% glycerol | 25 mM Tris, pH 7.5, 200 mM NaCl, 0.8 μM aprotinin, 2 μg/ml leupeptin, and 2 μM pepstatin A |
| **Cell lysis** | - | High-pressure cell disruption | Sonication | High-pressure Homogenizer | Sonication | Sonication |
| **Detergent for solubilization prepared in cell lysis buffer** | 1% (w/v) DMNG and 0.1% (w/v) CHS | 2% (w/v) DDM and 0.2% (w/v) CHS | 1.7% (w/v) LMNG or digitonin | 1% (w/v) LMNG and 0.1% (w/v) CHS | 1 % (w/v) DDM and 0.2% (w/v) CHS | 1% (w/v) Digitonin |
| **Affinity resin** | Tandem anti-FLAG & Ni-NTA | Anti-FLAG | Anti-FLAG | Anti-FLAG | Strep-Tactin | Strep-Tactin |
| **Wash buffer** | 0.01% GDN in cell lysis buffer | 0.02% (w/v) LMNG 0.002% (w/v) CHS in cell lysis buffer, plus 1 mM EDTA | 0.01% LMNG or 0.01% GDN 20 mM Tris-HCl, pH 7.0, and 150 mM NaCl | 40 mM Tris-HCl, pH 7.5, 150 mM NaCl, 10% glycerol (v/v), 1 mM DTT, 0.5 mM ATP-$Mg^{+2}$, 0.02% (w/v) GDN | 0.01% GDN, 25 mM HEPES, pH 7.5, 250 mM NaCl, 5% glycerol | 0.5% (w/v) Digitonin, 25 mM Tris, pH 7.5, and 200 mM NaCl |
| **Elution buffer** | 0.01% GDN in cell lysis buffer, plus 300 ug/ml FLAG peptide (for Anti-FLAG) or 300 mM imidazole for Ni-NTA) | 0.01% (w/v) LMNG 0.001% (w/v) CHS in cell lysis buffer, plus 500 ug/ml FLAG peptide, 50 mM HEPES, | 0.01% LMNG or 0.01% GDN 20 mM MES-NaOH, pH 6.0, 150 mM NaCl, and 5 M LiCl | 40 mM Tris-HCl, pH 7.5, 150 mM NaCl, 10% glycerol (v/v), 1 mM DTT, 0.02% (w/v) GDN, | 5 mM Desthiobiotin in wash buffer | 5 mM Desthiobiotin in wash buffer |

| | | | | | | |
|---|---|---|---|---|---|---|
| | | pH 7.5, and 300 mM NaCl | | plus 200 ug/ml FLAG peptide | | |
| **SEC buffer** | 0.006% (w/v) GDN, 25 mM HEPES, pH 7.4, and 150 mM NaCl | 0.01% (w/v) LMNG 0.001% (w/v) CHS, 25 mM HEPES, pH 7.5, 150 mM NaCl, and 0.5 mM EDTA | 0.01% LMNG or 0.01% GDN 20 mM MES-NaOH, pH 6.0, and 150 mM NaCl | 40 mM Tris-HCl, pH 7.5, 150 mM NaCl, and 0.02% GDN | 0.005% GDN, 25 mM HEPES, pH 7.5, 100 mM NaCl. For pH 5.5, 25 mM MES was used instead of HEPES | 0.5% (w/v) Digitonin, 25 mM Tris, pH 7.5, and 200 mM NaCl |
| **Protein concentration** | 13 mg/ml | 3.5 mg/ml | 2.5-5.0 mg/ml | 17 mg/ml | 10 mg/ml | 1 mg/ml |
| | | **Cryo-EM sample** | | | | |
| **Grid type** | Quantifoil Au 300 mesh, R1.2/1.3 | Quantifoil Cu 300 mesh, R0.6/1.0 or Nanodim R1.2/1.3 amorphous nickel titanium alloy (ANTA) grid | Quantifoil Cu 300 mesh, R1.2/1.3 | Quantifoil Cu 300 mesh, R1.2/1.3 | Ni-Ti Au 300 mesh (Nanodim Tech). | UltrAuFoil Au 300 mesh, R1.2/1.3 |
| **Pretreatments** | Grids glow-discharged | Grids glow-discharged and treated with 0.1% poly L-lysine hydrobromide | Grids glow-discharged | Grids glow-discharged | Grids glow-discharged | Grids glow-discharged, and 100 µM FOM added to protein sample |
| **Blotting** | 3 s, 8 ºC, 100% humidity | 2.5-3 s, 8 ºC, 100% humidity | 4 s, 6 ºC, 100% humidity | 4s, 6 ºC, 100% humidity | 3s, 4 ºC, 100% humidity | 2.5 s, 18 ºC, 100% humidity |
| | | **Cryo-EM data collection** | | | | |
| **Microscope** | Titan Krios (K3, 300 kv) | Titan Krios (K2/K3, 300 kv) | Titan Krios (K3, 300 kv), CDS mode | Titan Krios (K3, 300 kv) | Titan Krios (K3, 300 kv) | Titan Krios (K3, 300 kv) |
| **Pixel size** | 1.08 Å/px | 0.82 Å/px | 0.83 Å/px | 0.535 Å/px | 0.85 Å/px | 0.85 Å/px |
| **Total dose** | 50 $e^-/Å^2$ | 60 $e^-/Å^2$ | 66 $e^-/Å^2$ | 50-60 $e^-/Å^2$ | 52 $e^-/Å^2$ | 50 $e^-/Å^2$ |
| **Defocus range** | -1.3 to -1.8 um | -1.2 to -1.8 um | - | -1.5 to -2.0 um | -1.0 to -1.5 um | -1.0 to -2.5 um |

### Table S2: Comparison of the Cα RMSDs of hSIDT1-GFP (8V38) superposition on the existing ChUP family structures

| | 8JUL | 8JUN | 8HIIP | 8HKE | 7Y63 | 7Y68 | 7Y69 | 8WOT | 8WOR |
|---|---|---|---|---|---|---|---|---|---|
| **ECD dimer Cα RMSD** | 1.2 Å | 1.2 Å | 6.7 Å | 7.0 Å | 2.2 Å | 1.5 Å | 1.4 Å | 0.9 Å | 0.9 Å |
| **TMD dimer Cα RMSD** | 4.2 Å | 2.9 Å | 6.0 Å | 5.7 Å | 3.6 Å | 3.4 Å | 3.5 Å | 2.9 Å | 3.8 Å |
| **Full dimer Cα RMSD** | 4.1 Å | 3.1 Å | 8.2 Å | 9.0 Å | 3.3 Å | 3.3 Å | 3.3 Å | 2.6 Å | 3.3 Å |
| **ECD chain A Cα RMSD** | 1.2 Å | 1.2 Å | 6.7 Å | 9.9 Å | 2.2 Å | 1.5 Å | 1.5 Å | 0.9 Å | 0.9 Å |
| **TMD chain A Cα RMSD** | 4.2 Å | 2.9 Å | 6.0 Å | 5.7 Å | 3.6 Å | 3.4 Å | 3.4 Å | 2.8 Å | 3.9 Å |
| **Full chain A Cα RMSD** | 4.1 Å | 3.2 Å | 8.2 Å | 9.0 Å | 3.3 Å | 3.3 Å | 3.3 Å | 2.6 Å | 3.4 Å |
